# Improving viral protein clustering using both diversified protein profiles and structural information

**DOI:** 10.64898/2026.05.26.727815

**Authors:** Quentin Nugier, George Bouras, Clovis Galiez, Marie-Agnès Petit, François Enault

**Affiliations:** Université Clermont Auvergne, CNRS, LMGE, 63000 Clermont-Ferrand, France; Adelaide Medical School, Faculty of Health and Medical Sciences, The University of Adelaide, Adelaide, Australia; The Department of Surgery – Otolaryngology Head and Neck Surgery, Central Adelaide Local Health Network, Adelaide, Australia; Université Grenoble Alpes, CNRS, Grenoble INP, LJK, 38000 Grenoble, France; Université Paris-Saclay, INRAE, AgroParisTech, Micalis departement, 78350 Jouy-en-Josas, France

**Author notes:** corresponding author : François Enault.

## Abstract

Viruses are abundant, ancestral and potentially fast-evolving biological entities. As a result, their encoded proteins are diverse and identifying homologous relationships between sequences is as important for phylogeny and functional annotation as it is challenging. Traditional methods group viral proteins by sequence similarity, build HMM profiles for each protein family, and cluster further via profile comparisons. Here, we present an improved framework where HMM sensitivity is boosted by enriching reference virus HMM profiles with tens of millions of metagenomic sequences. This increases diversity within most protein families, raising the diversity index from less than 2 for 92.7% of clusters to a median value of 6. This enrichment of the profiles more than triples the number of homologies detected compared to the raw profiles. First-step clusters are then grouped more effectively using these relationships and further unified via structural predictions and comparisons. The sequence-enrichment strategy excels at linking small proteins, while structures better connect highly structured ones like tail and head proteins. Applied to 1.42 million proteins, our method yields 56,560 families—far fewer than 200,018 (sequence-based) or 135,048 (raw HMM)—revealing that prior approaches vastly overestimated viral protein diversity. The strategy of enriching the diversity of sequences of interest with external sequences, combined with the complementary use of structural information, highlights deep evolutionary links, offering a more accurate picture of viral protein evolution.

## Introduction

As all living organisms are infected by numerous viruses, viruses are present wherever there is life and in very large abundance (Mushegian 2020). Their diversity is great with an estimate of between 10 million and 1 billion different virus species (Koonin et al. 2022). This genomic diversity is also reflected in the proteins these viruses encode. The pangenome of various families of cultivated viruses is open (Rodrigues et al. 2022; Wang et al. 2020). Similarly, over 240,000 protein families in viruses populating marine ecosystems were found in metagenomic data (Zayed et al. 2021) and viruses on Earth were predicted to contain up to 10^10^ different protein families (Koonin et al. 2023). The clustering of viral proteins into homologous groups is a major challenge and a necessity to improve protein functional annotation and to better estimate the genes shared between viruses, which in turn will open the way for quantifying viruses functional and evolutionary distances and organize their taxonomy. Numerous databases of viral protein families, all based on sequence-based homology detection, have been developed over the last 20 years, such as POG, pVOG or EggNOG (Kristensen et al. 2013; Grazziotin et al. 2017; Hernández-Plaza et al. 2023). However, detecting all homology relationships between proteins remains a particularly difficult task for viral proteins. Not only is the number of different protein families large, but the internal sequence-level heterogeneity within most families is vast (Terzian et al. 2021). This internal diversity of protein families is often described as a direct consequence of the specific characteristics of viruses, which enable them to explore the protein sequence space extensively (Stern and Andino 2016; Koonin et al. 2022). It could also be due to specificities of part of protein families, rather than of viruses in general.

Whatever the cause, the great heterogeneity of viral sequences makes it impossible to identify all homology relationships using sequence similarity searches carried out by BLASTp-like tools (Blast: Altschul et al. 1990; Diamond: Buchfink et al. 2021; MMseqs2: Steinegger and Söding 2017). Accordingly, a commonly used strategy is to use a two-step clustering strategy: (i) group closely related proteins based on BLAST-like tool results, and (ii) build mathematical models based on intra-family sequence conservation and divergence for each protein cluster and compare these models. These models, such as sequence profiles based on position-specific scoring matrices (PSSMs) or Hidden Markov Models (HMMs), are built on multiple sequence alignments, encompassing evolutionary information which is lacking in single sequences. Thereby these models allow the identification of more distant homology. Such a two-step process has been recently implemented and used in several tools (VirClust: Moraru 2023; PhaMMseqs: Gauthier et al. 2022) and databases (VOGDB: Trgovec-Greif et al. 2024; PHROGs: Terzian et al. 2021; EnVhogDB: Pérez-Bucio et al. 2025).

Yet, the sensitivity of a given HMM depends entirely on the sequences it contains: a higher diversity of homologous sequences brings a higher sensitivity to the model. As the groups resulting from the second clustering (i.e. profile-profile) are, by nature, larger and more diversified than those from the first clustering, applying a third layer of comparison between them enables the detection of new homologous relationships. Such a strategy, based on a third clustering step, has been used in various studies (Olo Ndela et al. 2023, 2021) and corresponds to the so-called PSSCs (Protein-Super-Super-Clusters) of the VIRCLUST software (Moraru 2023). Nevertheless, this approach still depends on the diversity of the initial sequences considered. For viral genomes that are evolutionarily distant from other viruses, the proteins they encode will have few easily identifiable homologs, and their corresponding HMMs will not be very diverse, making it impossible to detect a greater number of homologs than what BLASTp-type tools have already done. As a result, we believe that many existing homology relationships remain undetected.

To propose a better organization into protein families for complete viral genomes, we used here external sequences, namely the tens of millions of viral environmental sequences available, to help diversify the HMMs of proteins from reference viruses and enable better clustering (Fig. 1). We used EnVhogDB (Pérez-Bucio et al. 2025), containing 53.6 million proteins organized into 2.2 million protein families, as it represents the most comprehensive database of metagenomic sequences. We (i) collected complete reference genomes, (ii) clustered their proteins, these first step clusters being termed VISEQs (for ‘VIral SEQuence group’), (iii) enriched each VISEQ by adding homologs from EnVhogDB, (iv) compared these enriched VISEQs and clustered them into clusters named VIRENs (‘VIral Remote groups built using sequence data ENriched with Environmental sequences’). Structure information has recently been shown to be able to capture more remote homology relationships between proteins (Barrio-Hernandez et al. 2023; Trgovec-Greif et al. 2024). We consequently (i) predicted the structure of a representative protein for each VISEQ, (ii) compared the similarities detected between these structures with those detected by sequence alignment strategies (iii) used these structure similarities as a third layer, to further group VIRENs into VIFOLDs (‘VIral clusters using FOLDing information’).

**Figure 1.**
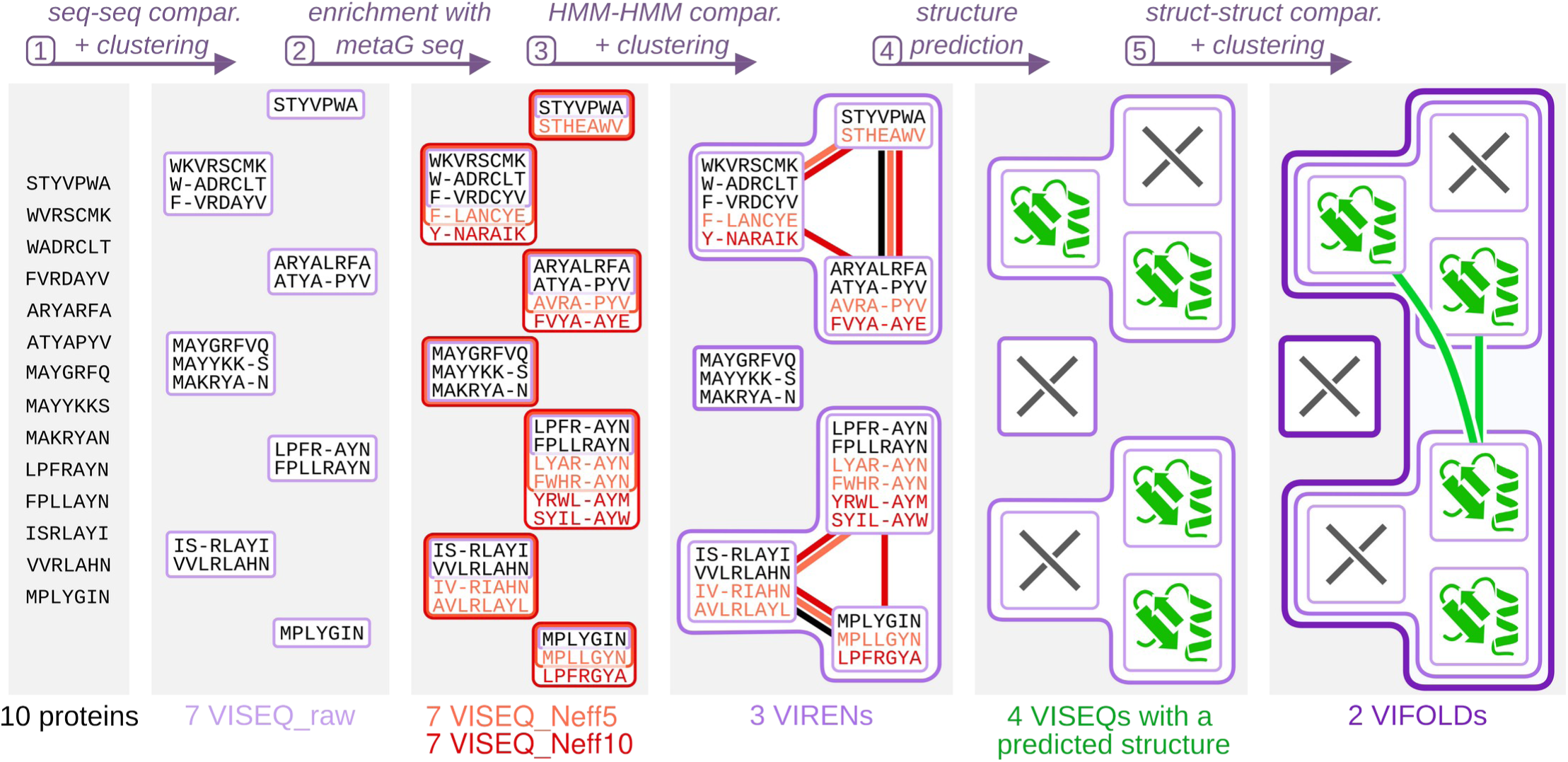
Overview of the viral protein clustering pipeline. As indicated at the top of the figure, the steps of the pipeline are: (1) reference viral proteins (RefSeq, INPHARED, etc.) are grouped into raw VISEQs based on sequence comparisons, (2) these VISEQs are enriched with metagenomic sequences from EnVhogDB with thresholds of Neff < 5 (orange, VISEQ_Neff5) and Neff < 10 (red, VISEQ_Neff10), (3) HMM-HMM comparisons between VISEQ_raw (black links), between VISEQ_Neff5 (orange links) and between VISEQ_Neff10 (red links) are used to group the VISEQs into VIRENs, (4) structures are predicted for the representative sequence of each VISEQ, (5) the VIRENs are grouped into VIFOLDs based on structure-structure comparisons. As shown at the bottom of the figure, the 10 proteins used as examples are grouped into 7 VISEQs, whose HMMs are enriched, then into 3 VIRENs, which are finally grouped into 2 VIFOLDs based on the similarities between the 4 VISEQs with a predicted structure.

## Results

### Description of the protein dataset and the first step clusters: the VISEQs

From the initial 75,195 genomic sequences, a subset of 36,172 genomes (40,728 genomic sequences) was kept after the removal of redundant genomes and almost identical genomes (>98 % identity and >95 % coverage). From these sequences, 1,524,198 proteins were obtained : 556,453 proteins from RefSeqVirus (17,312 genomic sequences), 556,307 from INPHARED (13,921), 256,996 from PHROG (6,329) and 112,984, 40,752 and 706 for additional transposable phages (2,196), giant viruses (833) and microvirus (137) respectively. Protein sequences were then compared with MMseqs2 (coverage>80%) and clustered into 336,106 VISEQs, made of 123,326 clusters of at least two proteins and 212,780 singletons. HMMs were built for all of these VISEQs and were named VISEQ_raw.

### Diversifying VISEQs with external environmental viral proteins

HMM from these VISEQ_raw had low Neff values (median 1.0; maximum 8.1), all singletons having by definition a Neff value of 1 (Fig. 2). Concerning the 123,326 non-singleton VISEQs with at least 2 proteins (designated as clusters on Fig. 2), 80.2 % (98,928) had a Neff lower than 2 and 94.7 % (116,841) had a Neff lower than 3, confirming our initial hypothesis that first step VISEQ clusters have a low diversity. Each of these 336,106 raw HMMs were then enriched using the approximately 2.2 M HMMs of EnVhogDB. As a result, the diversity of the VISEQs was greatly increased (Neff median: 6.0; maximum: 14.8). Considering the 123,326 VISEQs of at least two proteins, as many as 82.1 % of VISEQ HMMs had Neff values increased by a factor of 2 at least, and only 4.3 % were not enriched at all. Importantly, for the 212,780 VISEQs composed of only one sequence, *i.e.* singletons, more than 82.0 % of them had homologs in EnVhogDB (Neff > 1 after enrichment, Fig. 2). Diversification allowed a drastic reduction of the proportion of viral protein clusters with a low diversity (Neff value<2): they initially represented 92.7 % (311,708) of all VISEQs, and only 19.5 % (65,547) after the enrichment step. At the other extreme, 12.4 % of HMMs had their diversity increased drastically, their Neff value exceeding 10. As mentioned in the HHblits software recommendations, the specificity of subsequent similarity searches may be diminished for such diverse HMMs. Thus, all 336,106 HMMs had their diversity limited to a Neff value of 10, resulting VISEQ_Neff10. In addition, a version of the HMMs was also computed with a Neff value limited to five, producing the VISEQ_Neff5 set.

**Figure 2.**
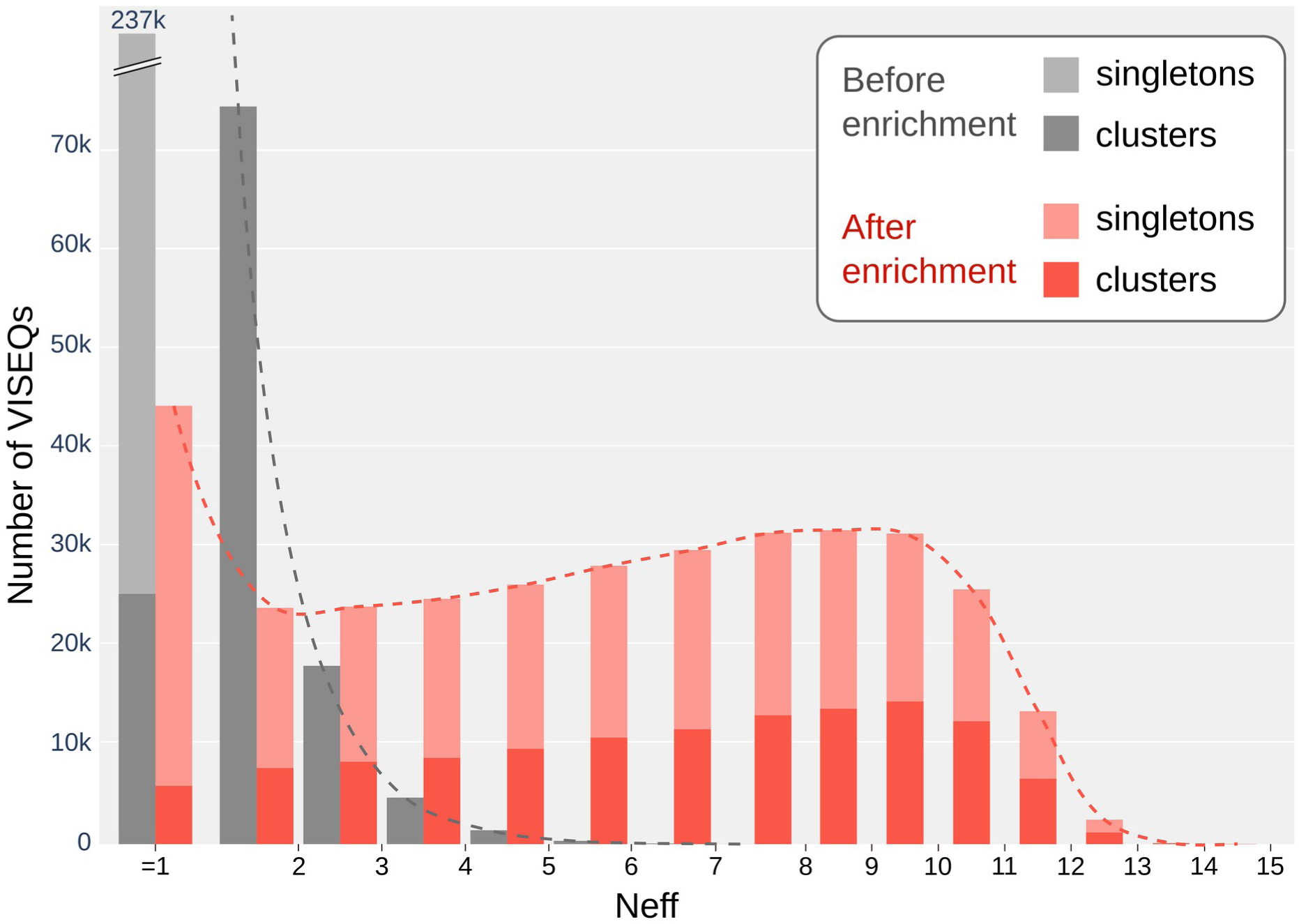
Histograms of Neff values for VISEQs before (grey) and after (red) enrichment of their diversity with EnVhogDB sequences. VISEQs containing only one protein are termed singletons whereas VISEQs with at least two proteins are named clusters. Singletons and clusters made of identical proteins have by definition a Neff of 1 before enrichment (grey bar on the left, for a total of 237 K VISEQs), and 13.1 % of VISEQs were not enriched by any EnVhogDB sequence (red bar on the left).

### Building VIRENs by comparing and clustering VISEQs

The three sets of HMMs (VISEQ_raw, VISEQ_Neff5 and VISEQ_Neff10) were compared to themselves using one or two iterations of HHblits. We calculated the Fisher’s index between pairs of VISEQs, combining the similarities detected by the five resulting methods. By retaining only those links with an index of at least 50, we are able to exclude the least reliable links between VISEQs, i.e. those identified by only one of the five methods and/or those with only high evalues for one or more of the methods. For the 3.06 M pairs of VISEQs retained, the five methods contribute to varying degrees, accounting for between 0.31 and 2.71 million significant similarities between two VISEQs (Fig. 3; Fig. S1 for further details). Starting from the VISEQ_raw profiles, their comparison with one iteration, a method similar to the one used to build PHROGs, identified only 10 % of the final pairs uncovered across all methods (0.31 M matches). This value increased to 30 % when using VISEQ_raw and two similarity search iterations. On the contrary, starting from the enriched VISEQs (VISEQ_Neff5 or Neff10) allowed to reveal 99 % of all links, almost all similarities detected between VISEQ_raw being also identified with enriched HMMs (Fig. 3A). Each of the five comparison methods used provided a specific contribution (Fig. 3B), the intersection of all methods being small (0.26 M links) mostly due to the low number of hits found between VISEQ_raw with one iteration. We conclude that enriching VISEQ_raw profiles using EnVhogDB allows a vast increase in sensitivity.

**Figure 3.**
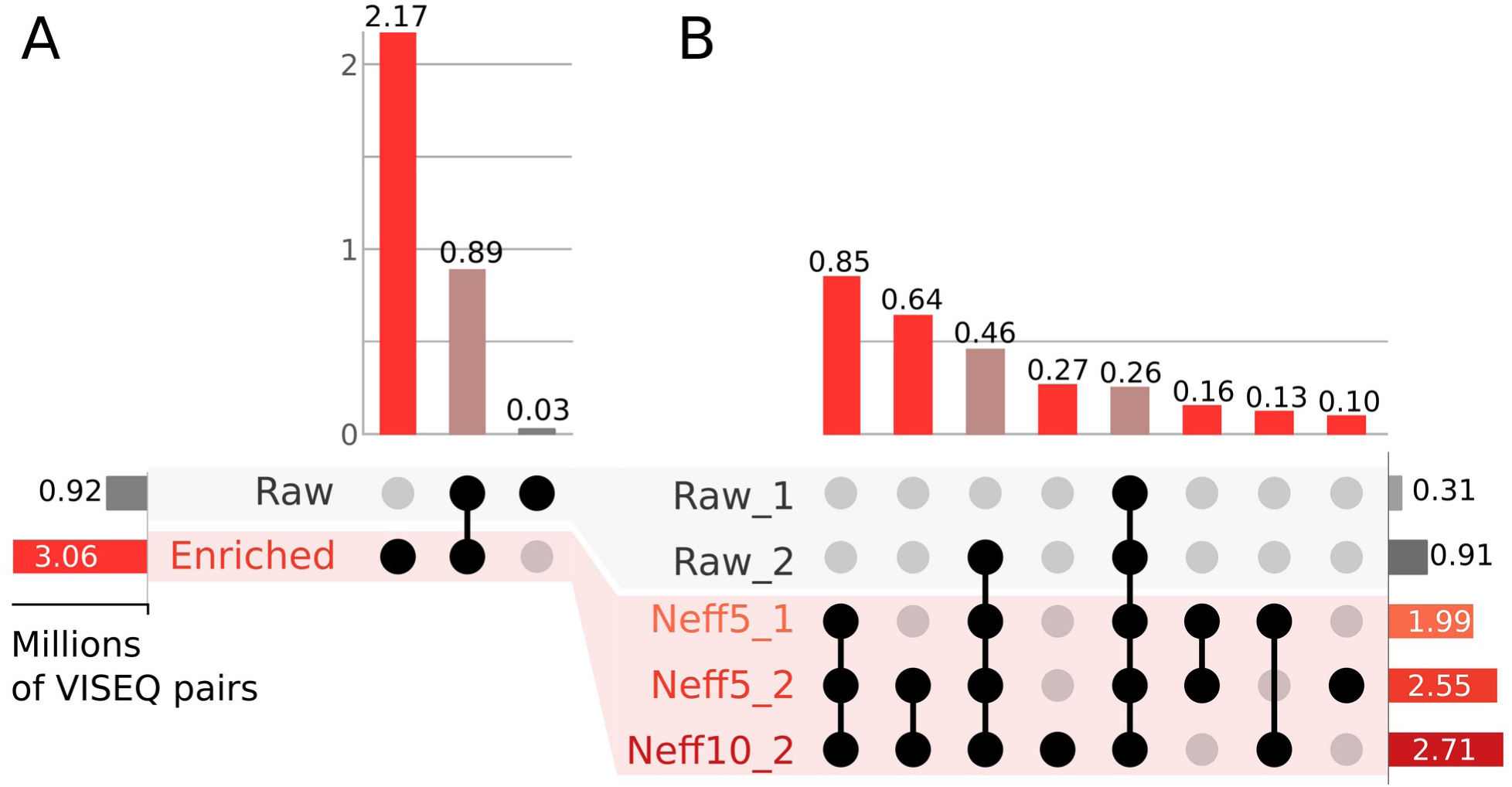
Number of links detected between VISEQs using three sets of HMMs (Raw, Neff5 and Neff10) compared using either one or two iterations (suffix 1 or 2), considering only the similarities between the 3.06 M pairs of VISEQs finally retained (Fisher index of at least 50). (*A*) Similarities detected between VISEQ_raw, using one and two iterations, were gathered in the grey ‘Raw’ line, the red ‘Enriched’ line gathering similarities obtained with Neff5 (one and two iterations) and Neff10 (two iterations). Similarities detected by both raw and enriched HMMs are displayed as pinkish-brown. (*B*) Detailed numbers for the five different strategies, only combinations representing more than 0.1 M links are shown.

Using these 3.06 M links between VISEQ pairs, the 336 K VISEQs were clustered into 164,276 VIRENs. Considering the initial 1.52 M proteins, 1.42 M were gathered into 64,185 VIRENs with more than one protein (average of 22.2 proteins, the largest VIREN containing 14,655 proteins) and only 100,091 were left as singletons.

### Predicting VISEQs structures and using them to cluster VIRENs into VIFOLDs

To increase sensitivity, structure predictions and comparisons were performed at the VISEQ level, rather than the VIREN level. Using AlphaFold2, the structure was predicted for all 336 K VISEQ representative sequences and 194,000 of them (58 %) passed the minimum confidence filters. These retained structure predictions were distributed inside 76 K distinct VIRENs, with a maximum of 1,470 structures inside a single VIREN. These 194 K structures were then compared to each other using Foldseek and Foldseek-TM (evalue < 0.01, TM-score > 0.5), and 1,64 M structural similarities were detected, involving 114,666 VISEQs.

Among these structural links, 110,852 connected different VIRENs, and in total 25,837 VIRENs were candidates to the higher-level clustering. To group only those VIRENs whose VISEQs are well connected, a minimal number of links, corresponding to half of the geometric average of potential links was defined (Fig. S2 for a detailed example). It corresponds to the best compromise between a small number of final clusters and a high quality of links between VIRENs inside a same VIFOLD (Fig. S3). Filtering links between VIRENs using this threshold, only 73,318 links between 19,902 VIRENs were retained. As a comparison, using only the structure prediction of the ‘central’ protein of each VIREN as in VOGDB, this central protein being the one having the higher sum of sequence alignment scores with the other protein of the same VIREN, only 36,689 matches between 14,422 VIRENs were detected. Using these 73,318 links, 149,973 clusters were created and named VIFOLDs. Some VIFOLDs only have one VIREN, either because the VIREN has no predicted structure (due to the predicted structure for constituent VISEQs not passing confidence filters) or because no (or not enough) homologous structure was detected in other VIRENs. The largest VIFOLD was composed of 440 VIRENs (vifold_43). Considering the initial 1.52 M proteins, 1.42 M were gathered into 56,580 VIFOLDs with more than one protein (avg of 25.1 proteins, the largest VIFOLD containing 19,651 proteins) and only 93,392 were left as singletons (Fig. S4).

### Comparing the clusters to homologous groups of PHROG and VOGDB

The results of our three clustering steps were then compared with the viral families established by other databases: (i) the PHROGs, created using a two-step clustering method corresponding to our method Raw_1 and (ii) the VOGs, VFAMs and VFOLDs of the recently proposed VOGDB (not to be confounded with pVOGs, doi: 10.1093/nar/gkw975), built using a three-step clustering similar to our strategy, but with important departures. Indeed, the first-layer VOG clusters were made with PSI-BLAST followed by a COG triangles clustering method, instead of MMseqs2 alignments followed by MCL clustering. The second-layer clusters, VFAMs, was similarly based on HMM-HMM comparisons and clustering, but with no enrichment step using external sequences. Finally, the predicted structure of VFAM representative sequences were compared and clustered into VFOLDs using Foldseek, whereas we predicted and clustered the structures of the first-level clusters, the VISEQs, which contain far more sequences.

To compare the performance of the different methods, proteins common to all three databases were identified along with their cluster number in each of the seven datasets (VISEQs, VIRENs, VIFOLDs, PHROGs, VOGs, VFAMs, VFOLDs). Altogether, half of the 640 K proteins were grouped into clusters containing at least 53, 108, 116, 136, 170, 222 and 303 proteins for VISEQs, VOGs, PHROGs, VFAMs, VFOLDs, VIRENs and VIFOLDs respectively (these values correspond to those shown with the horizontal bar in Fig. 4). Unsurprisingly, VISEQs were very small clusters compared to other methods due to the high coverage threshold of 80 %. As a result of the small size of VISEQs and of their limited internal diversity (Fig. 2), PHROGs that are based on the comparisons of these clusters were the second smallest groups. VOGs were significantly larger than these two clustering, due to the use of PSI-BLAST, allowing to incorporate the diversity of all similar sequences, with no coverage restriction. The comparisons and clustering of these VOG HMMs generated only slightly larger VFAMs, but the comparisons of the structure prediction of their representative sequence generated larger VFOLDs. Interestingly, VIRENs were as large as these VFOLDs from VOGDB, even though built using sequence information alone. This highlights the impact of using external sequences to diversify groups in order to detect more distant similarities. Finally, further clustering of these VIRENs into VIFOLDs allowed us to build the largest clusters.

**Figure 4.**
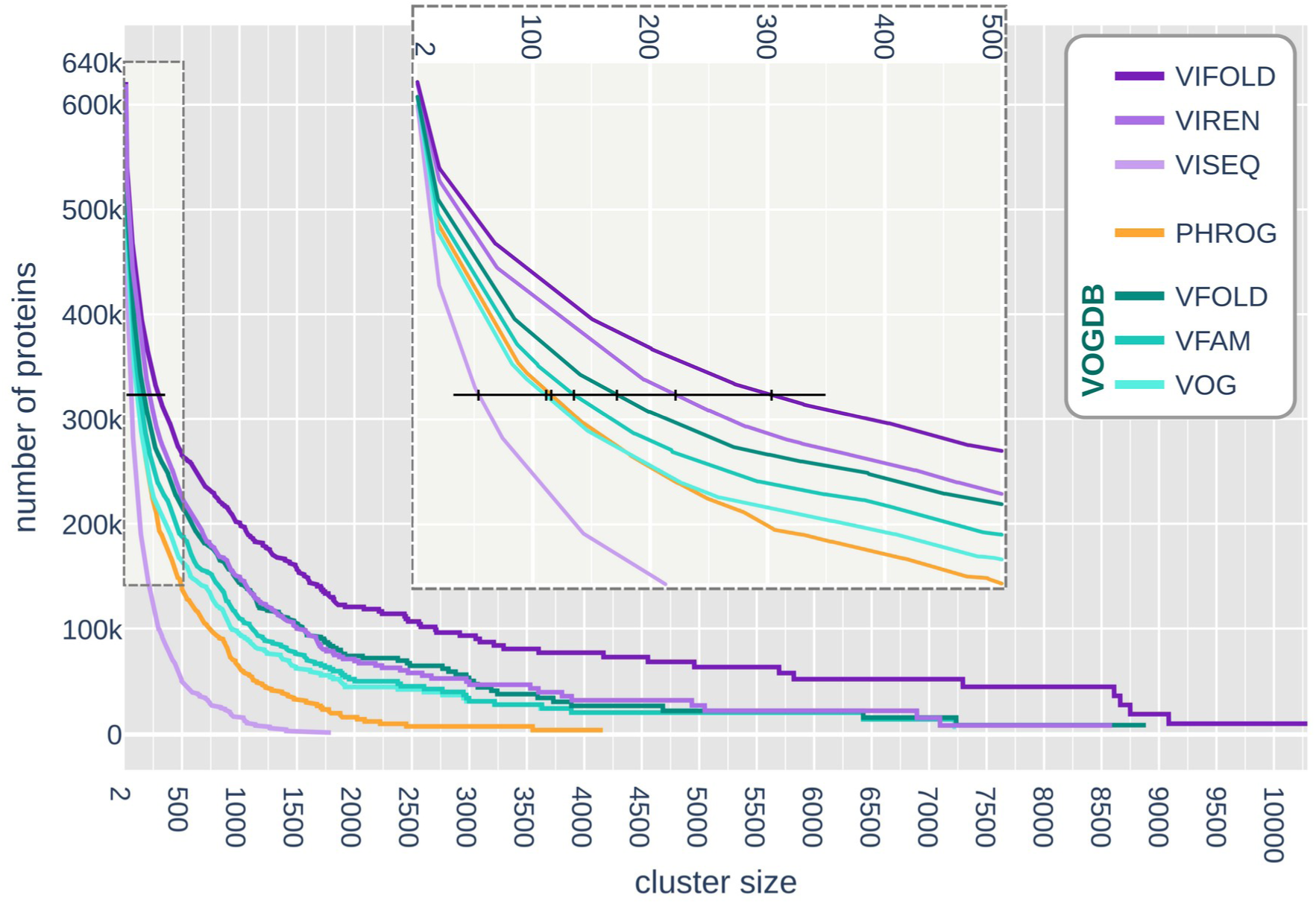
Cumulative number, starting from the right, of clustered proteins for our three clustering steps (shown in shades of purple), for PHROG (in orange) and for the three VOGDB clustering steps (in shades of turquoise), taking into account only the 640,096 proteins common to these three databases. The largest cluster for each method corresponds to the right-hand end of the curves, and these contain 1795, 4161, 8592, 7233, 7236, 8886 and 10291 proteins for VISEQ, PHROG, VIREN, VOG, VFAM, VFOLD and VIFOLD, respectively. As can be seen in the enlarged view (at the top center) of the left-hand side of the curves, half of the 640 K proteins (represented by the black horizontal bar) are in clusters containing at least 53, 108, 116, 136, 170, 222 and 303 proteins for VISEQ, VOG, PHROG, VFAM, VFOLD, VIREN and VIFOLD respectively.

In addition, clustering the whole 1.52M protein dataset using a two-step method as in PHROG or VIRCLUST or a still widely adopted BLASTp-like approach (cov>50%) gives an overview of how both of these strategies are misfitted for remote homology detection and downstream building of homologous protein groups. While 56,560 VIFOLDs were built, 84,070 PHROG-like clusters and 112,672 sequence-based clusters of at least two proteins were produced. Furthermore, the present clustering left only 93,392 unclustered singleton proteins, compared to 144,370 and 180,738 singletons for the other two approaches respectively (Fig. S5).

### Exploring the differences between similarities detected by HMMs and structures

To further understand the strengths and weaknesses of the sequence and structural comparison methods, the 3.06 M HMM links and 1.64 M structural links between VISEQs were compared. First, only 0.59 M links were detected by both methods. This is due, for a large part, to the 42.3 % VISEQs with no structure predicted. We thus limited the analysis to the 194 K VISEQs with a predicted structure, and found interestingly only a partial overlap between the two approaches, none showing a clear superior performance. Indeed, the same 0.59 M links were both detected by HMM-HMM comparisons (2.18 M links) and structure-structure comparisons (1.64 M links).

For both sequence and structural comparison methods, protein length is the main factor. For structure-based analyses, protein length has an impact on the number of similarities found as (i) the structure of very short (<50 AA) and long proteins (>1000 AA) is less frequently confidently predicted (Fig. S6) and (ii), even when considering only VISEQs with confidently predicted structures, similarities were harder to detect for proteins smaller than 250 residues and for proteins larger than 800 residues (Fig. S7). Considering sequence information, the sensitivity of HMM comparisons is only lower for very short or very long proteins (<50 residues or >800 residues, Fig. S7). As a result, many more similarities were detected by HMM than by structure comparisons for proteins smaller than 150 residues. When considering only annotated VISEQs, 250,088 links between two VISEQs were detected using HMMs and 91,100 using structures (Fig. 5B).

**Figure 5.**
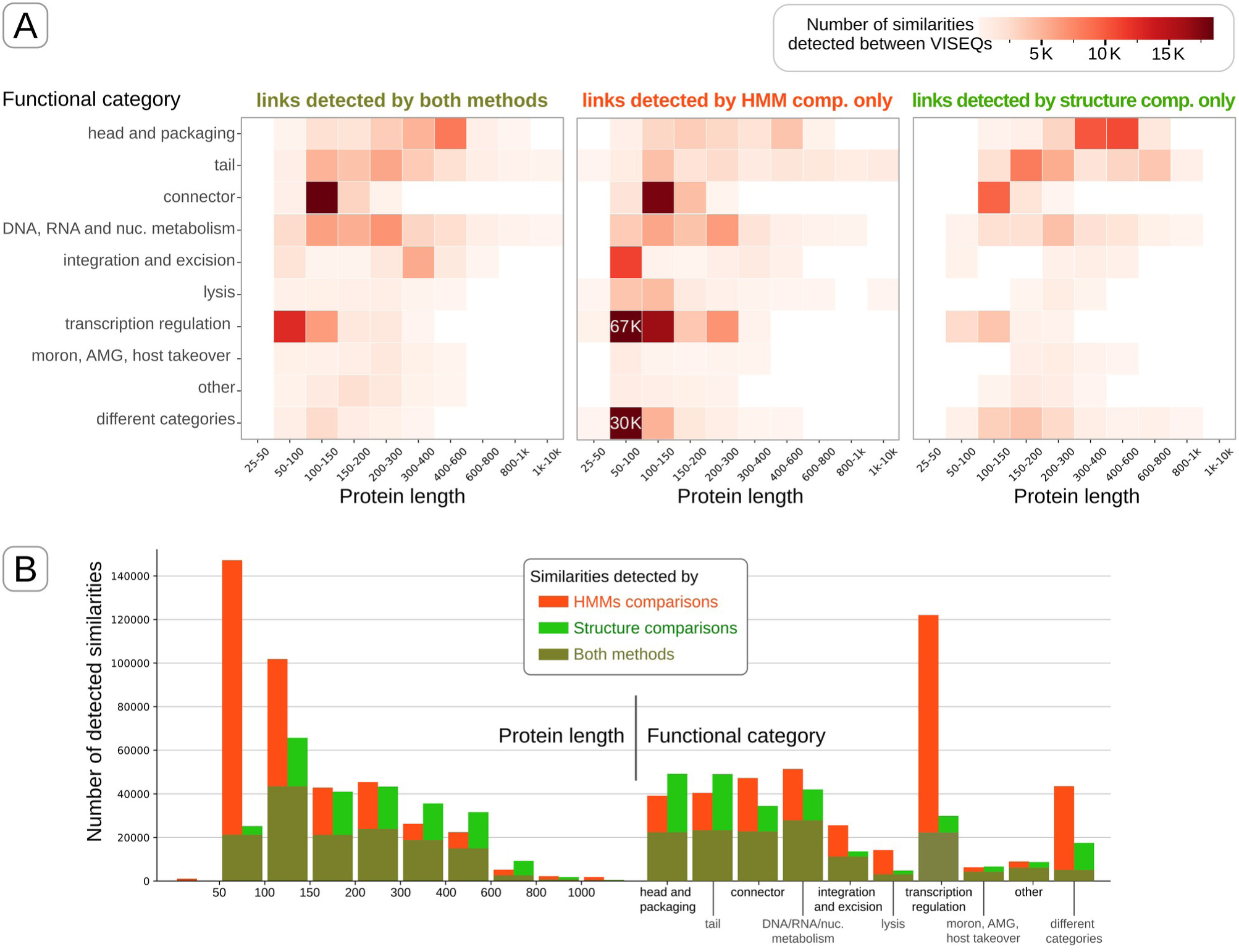
Analysis of the similarities between VISEQs detected by HMM-HMM and structure-structure comparisons. Only VISEQs with an annotation are here considered. (*A*) Heatmaps displaying the number of similarities detected by both methods and specific to each method. For a clearer colour scale, two very large values are directly written in two cells (67 K for 67,166 and 30 K for 29,963 links). (*B*) The same results are displayed in the form of two histograms, categories being protein length intervals and functional categories.

We then sought to determine whether the links between VISEQs across different functional categories were more frequently revealed by structural comparisons or by HMM comparisons. Interestingly, most links detected solely by structure comparison concerned proteins in the ‘Head and packaging’ and ‘Tail’ categories (Fig. 5B), *i.e.* those involved in virion formation, especially for proteins longer than 150 residues (Fig. 5A). The links detected only by HMMs concerned several other categories, namely ‘DNA, RNA and nucleotide metabolism’, ‘Transcription regulation’, ‘Integration and excision’, ‘Lysis’, and ‘Connector’ (Fig. 5B), especially for proteins smaller than 150 residues (Fig. 5A).

We also noted a high number of links between proteins annotated in different functional categories (55,524 links), and a majority of them concerned short proteins connected by HMMs (30,203 links). Most of the discrepancies were due to the annotation itself rather than to VISEQ pairs that had been incorrectly identified as similar. Indeed, more than half of these pairs consisted of small proteins (<100 residues) with a DNA-binding domain, with the VISEQs annotated either as DNA-binding proteins (belonging to the ‘DNA, RNA and nucleotide metabolism’ category), or as transcription regulators (belonging to the ‘Transcription regulation’ category), or as integrases/excisionases (belonging to the ‘Integration and excision’ category).

## Discussion

The goal of this study was to determine the best methodology for clustering viral proteins using current tools and data, both to produce a clustering of proteins of current reference viruses and to propose a methodology for researchers to establish such a protein clustering on new viral data. Our study has shown that enrichment with external sequences, such as the tens of millions of metagenomic sequences, facilitates the identification of evolutionary links between sequences of cultivated viruses. This enrichment of HMMs was combined with a structural prediction followed by a structure-structure comparison approach, that has now become feasible at a large scale. It enabled proteins to be grouped into larger clusters than those obtained using existing methods, even the method recently proposed in VOGDB. One consequence of this large clustering effort is that the number of singletons now represents only 6.1 % of all proteins included in this study. These singletons could correspond to proteins of undersampled viruses, but we rather suspect, as these proteins are not encoded by viruses that are particularly distant to others in evolutionary terms, that they are poor predictions of protein-coding genes, and/or truncated proteins due to IS insertions. Not considering these singletons, the 1,430,805 distinct proteins from all known viruses were grouped into only 56,580 VIFOLDs. This gives a very different picture, compared to the 200,018 clusters generated (for the same protein set) using standard methods based on BLASTp-like tools (cov>50%). BLASTp-based clustering methods are still used in the analysis of most new viral protein dataset, although we show here that it overestimates the number of protein families by a factor of three.

Enrichment by environmental sequences highlights the great internal heterogeneity of viral protein families, the diversity of most families being huge (median Neff of 6). This internal diversity of protein families is often described as a direct consequence of virus specific characteristics (lifestyle, large population number, rapid life cycle, absence of repair systems), which results in potential rapid mutagenesis and generation of numerous possible versions of the proteins (Stern and Andino 2016; Koonin et al. 2022). In addition to these general characteristics of viruses that affect all protein families, we believed that the diversity of the protein families that make up the virion could be more significant. Here, however, a great internal diversity was observed for all protein sizes and all functional categories: the structural proteins involved in virion formation, were indeed highly diverse, but so were the proteins involved in nucleic acid metabolism or transcription regulation, and in every other category (Tab. S8). It would be interesting to compare more precisely this diversity with that of cellular proteins in order to better understand and decipher the evolution and specific features of the viral world.

Even if proteins of all sizes and functional categories are diverse, we observed that the similarities captured by HMM-based and structure-based comparisons were not the same. First, short sequences were less suitable for structure comparisons, because fewer structures were predicted and because their similarities were more difficult to determine. This may simply be due to the fact that these proteins contain less structural information, but we cannot exclude that they are also intrinsically less structured due to disorder or conformational flexibility. Conversely, we found that links between proteins with very precise folds, such as capsid or tail proteins, were better identified by structure analysis, perhaps because these proteins are too diverse at the sequence level to be represented in the form of an HMM. This internal sequence heterogeneity difficult to capture by alignment methods may be explained by the type of constraints on their conformation and interaction with other proteins likely very different from those of enzymes, for example.

In addition to providing a better understanding of protein diversity and the best ways to identify evolutionary relationships, we believe that these protein families from reference viruses have real operational value. Indeed, even though environmental viral datasets consist of hundreds of millions of proteins and millions of protein families (Pérez-Bucio et al. 2025), the present dataset contains (almost) all of the proteins whose annotations have been determined experimentally. Moreover, as the HMM profiles of the families produced have been diversified by environmental sequences, the 56 K VIFOLDs also encompass most of the viral data known to date. Containing both all currently known functional information and sequence diversity, this dataset is therefore particularly interesting because its small size makes it easy and quick to compare with other data. In order to use these VIFOLD clusters to annotate proteins from new viruses, our next goal will be to harmonize, verify and curate the functional annotations of these clusters, which are currently generated automatically, in order to provide virologists with up-to-date and homogeneous viral protein annotations.

## Methods

### Creation of a set of reference viruses and their proteins

Complete virus genomes were retrieved: (i) reference virus genomes from RefSeqVirus (January 2024), (ii) phage genomes from GenBank identified as complete from INPHARED (April 2024), (iii) genomes from the PHROG database, mainly for provirus genomes, (iv) a selection of giant virus genomes curated by specialists (January 2024), (v) a set of complete transposable prophage genomes (Zhang et al. 2023; Olo Ndela et al. 2021) and (vi) complete circular genomes of microviruses assembled from metagenomics and selected to best represent the diversity of this family (F Enault, pers. comm.). As the sources used contain redundant genomes and almost identical genomes from closely related isolates, these were deduplicated by grouping very similar genomes based on their intergenomic similarity, calculated using the formula described in the VIRIDIC tool (Moraru et al. 2020). Genome pairs with more than 98% identity and more than 95% coverage were grouped together, and a single genome was retained for each of these groups, with preference given to genomes from RefSeqVirus. For genomes for which no proteins were available or for which the coding potential was abnormally low (sum of all gene lengths lower than 30% of the genome length), proteins were again predicted using Prodigal (version 2.6.3; Hyatt et al. 2010).

### Construction of the first layer of clusters: VISEQs

Protein clusters were generated using the *easy-cluster* command of MMseqs2 (version 15-6f452; coverage > 80%, evalue < 0.001), each cluster being called a VISEQ. For each VISEQ, a multiple alignment was computed with ClustalOmega (Sievers et al. 2011; version 1.2.2; 5 iterations). In order to remove positions corresponding to wrongly predicted start position or to rare insertions, these alignments were converted into the a3m format, a consensus sequence being added that only contains the positions with less than 50% of gaps, using the *hhconsensus* command with the -M 50 parameter. HMMs were then built from each alignment using *hhmake*, and were termed VISEQ_raw. These commands are part of the HH-suite (v. 3.3.0; Remmert et al. 2012). All initial proteins were compared to proteins of the PHROG database v5 using MMseqs2 (-s 9, cov>70%, id>50%, evalue < 0.001) and the best match against PHROG-annotated proteins was transferred. For VISEQs made of proteins annotated with different functional or category terms, an expert analysis was performed and the most appropriate annotation was selected. To avoid hits between imprecise structural proteins (such as ‘Virion structural protein’, by default in the ‘Head and packaging’ category but some of which are tail or neck proteins), a new category was created as ‘Undefined structural protein’ and excluded from the functional analysis. Finally, a representative protein was also determined for each VISEQ using the MMseqs2 *easy-cluster* command.

### Enriching VISEQs with external sequences

To increase their internal diversity, VISEQs were compared to the viral protein families of the EnVhogDB by HMM-HMM comparisons with the hhblits_omp command, a parallelized version of HHblits (probability ≥ 0.9; iter = 2). Sequences of EnVhogDB HMMs detected as similar were added to the VISEQ MSAs to increase their diversity. To assess the extent to which enrichment increases alignment diversity, the Neff of every VISEQ was calculated, before and after the enrichment step.

This diversity index is the average, over all columns of the MSA, of the Shannon diversity index, *i.e.* the number of different residues present per column, weighted by the proportion of each residue (Meier and Söding 2015). By enriching VISEQs with EnVhogDB sequences, HMMs are likely to accumulate a great deal of diversity. Incorporating too much diversity in each HMM can be counterproductive. Indeed, this diversity is taken into account in the calculation of the evalue determining the significance of the similarity between HMMs. Thus, similarities between two truly homologous HMMs, each with very high diversity, will be detected but will have large and non-significant evalue. To control this undesirable effect, HMMs had their diversity lowered down to a Neff value of 10 and 5 using the hhfilter tool, resulting in VISEQ_Neff10 and VISEQ_Neff5 respectively.

### Compare VISEQs in different ways and building the second-layer clusters: VIRENs

VISEQs were compared using HHblits (probability ≥ 0.9, coverage ≥ 0.6) using five different combinations of HMMs and parameters. First, VISEQ_raw were compared to themselves with one iteration. This strategy of comparing raw first-step clusters, termed here Raw_1, corresponds to the strategy used to build homologous groups of the PHROG database. VISEQ_raw were also compared to themselves using two comparison iterations, the resulting similarities being termed Raw_2. Likewise, VISEQ_Neff5 were compared to themselves with one or two iterations, forming the Neff5_1 and Neff5_2 approaches, respectively. In addition, VISEQ_Neff10 were ultimately used to detect similarities between very distant VISEQs, particularly to identify highly diverse VISEQs that are homologous to VISEQs with low diversity. Consequently, we identified similarities between VISEQ_Neff10 with only two iterations, producing a single set of Neff10_2 links.

For each pair of VISEQs A and B detected as similar by at least one of the five above methods, the p-values of the similarities between A and B (Raw_1, Raw_2, Neff5_1, Neff5_2 and Neff10) were mixed using the statistic of *Fisher’s combined test* (Fisher 1932), defined as the sum of the natural logarithm of all p-values, multiplied by -2. As the p-values are not independent, the resulting statistic was not used to perform the test but as a measure of similarity for all pairs of VISEQs, referred to as ‘Fisher index’. For example, the Fisher index for a pair of VISEQs with a single similarity with a p-value of 10^-10^ is 46.04, and the Fisher index for a pair of VISEQs with two similarities with p-values of 10^-6^ is 55.26. To cluster VISEQs into larger groups, only pairwise similarities with a Fisher index ≥ 50 were used. In addition, to prevent transitivity issues, only VISEQ pairs with at least one hit with a reciprocal coverage ≥ 70 % were considered. Based on the resulting similarities, VISEQs were then clustered using MCL (Van Dongen, 2008; -I 2.0, no edge weight). Hereafter, the resulting clusters are referred to as VIRENs.

### Structure predictions, comparisons and construction of the third-layer clusters: VIFOLDs

For all the representative proteins of VISEQs, structure prediction was conducted as follows. Firstly, only VISEQs with representative sequences of 3000 residues or below were considered, due to the computation VRAM limitations of the GPUs used for structure prediction, leaving 335,676 VISEQs for which we generated structure predictions. We then generated two MSAs for each VISEQ. The first MSA was using the default ColabFold (Mirdita et al. 2022) ‘colabfold_search’ parameters utilizing the default uniref2302_30 and colabfold_envdb_202108_db (ColabFold DB) databases. The second MSA was created by also adding the ‘colabfoldv’ viral database containing approximately 130M viral proteins to the two default colabfold_search databases (https://github.com/gbouras13/colabfoldv), which has been shown to result in richer MSAs and improved viral protein structure predictions compared to the default ColabFold databases (Bouras et al. 2025).

We then used AlphaFold2 (Jumper et al. 2021) implemented via Colabfold v1.5.5 to conduct protein structure predictions using AMD MI250x GPUs on Setonix at the Pawsey Supercomputing Research Centre. Specifically, we used a customised Singularity container available from https://quay.io/repository/sarahbeecroft9/colabfold/rocm6.0.0_cpuTF. AlphaFold2 was run in batch mode (colabfold_batch) with 1 model (‘--num-models 1’) with the default 3 recycles. No AMBER relaxation or templates were used. For each VISEQ, we selected the protein structure prediction with the highest mean pLDDT between the ColabFold default MSA and colabfoldv enriched MSA as the best model to proceed with. Short protein sequences may have a higher pLDDT, even in the case of spurious structure predictions (Monzon et al. 2022). To be considered of sufficient quality, the structures of proteins with fewer than 70 residues had here to have a pLDDT greater than 80, whereas a threshold of only 70 is required for those with more than 70 residues.

All predicted structures were then compared using Foldseek (van Kempen et al. 2024) and Foldseek-TM, an implementation of TM-align (Zhang and Skolnick 2005), two alignment methods integrated into the foldseek software (version 7.04e0ec8), with a minimum threshold of 0.6 for reciprocal coverage. Next, as Foldseek calculates an evalue whereas TM-align does not, pairs of structures showing a similarity with an evalue of less than 0.01 according to Foldseek were selected. For each selected pair of structures, the best TM-score from the two comparison methods (Foldseek and TM-align) was retained if the coverage was greater than 70%. Finally, all similarities between representative VISEQ protein structures with a TM-score of 0.5 or higher (Xu and Zhang 2010) were retained and considered as structural links.

VISEQ pairs belonging to different VIRENs, but for which structural similarities were found were then used to cluster VIRENs. To avoid cases in which poorly connected VIRENs are grouped, a threshold on the number of structural similarities between their respective VISEQs required to link two VIRENs was determined (Fig. S3). For two VIRENs having respectively N and M predicted structures, the maximal number of links between them is N*M, and their expected average number of links is sqrt(N*M). Considering half of this value as a threshold (L ≥ sqrt(N*M)/2, with L being the number of observed links between the two VIRENs), pairs of VIREN were used to cluster VIRENs using the MCL software, generating VIFOLDs, for ‘VIral FOLDs’.

### Comparison with other established databases and approaches

Proteins present both in the dataset described here and in PHROG and VOGDB were detected by comparing proteins of these two databases to the 1.52 M proteins using MMseqs easy-search (cov>90%, id>90%) best hits. For each protein identified in the three datasets, the cluster they belong to in all database of origin were retrieved. To compare the different clustering methods to standard clustering procedure, basic clusterization using a BLASTp-like tool was also done using MMseqs easy-cluster (cov>50%, evalue < 0.001, with --cluster-reassign) and a PHROG-like approach using Raw_1 links only (cov>60%, probability>90%).

### Data access

Scripts and information on the pipeline are available at https://gitlab.com/quentin_ngr/virendb.

All genomes, protein sequences and information, as well as MSAs, HMM profiles and predicted structures of protein clusters are downloadable at https://doi.org/10.57745/DNFV8P.

## Acknowledgements

This work was supported with the assistance of resources and services from Phoenix HPC at the University of Adelaide and Pawsey Supercomputing Research Centre, which is supported by the Australian Government. We would like to thank Fabien Voisin and Sarah Beecroft for their assistance in operating ColabFold at scale at Phoenix and Pawsey respectively, with extra acknowledgement to Sarah for containerising ColabFold for use on Setonix’s AMD GPUs. We are grateful to the INRAE MIGALE bioinformatics facility (MIGALE, INRAE, 2020. Migale bioinformatics Facility, doi: 10.15454/1.5572390655343293E12) for providing computing and storage resources. We are grateful to the Mésocentre Clermont Auvergne University for providing computing and storage resources. This work was supported in part by the European project Asclepius (Grant Agreement number 101159555).

## Author contributions

F.E. and M.A.P. conceived and led the study. Q.N. conducted data analyses. C.G. provided support for software usage and statistics. G.B. conducted structure predictions and comparisons. F.E. and Q.N. prepared the manuscript with input from all authors. All authors reviewed the manuscript.

